# Limits to the spread of an obligate social cheater in the social amoeba *Dictyostelium discoideum*

**DOI:** 10.1101/2023.01.17.524501

**Authors:** James Medina, David Queller, Joan Strassmann

## Abstract

Cooperation is widespread across life, but its existence can be threatened by exploitation. Social cheaters can be obligate, incapable of contributing to a necessary function, so that spread of the cheater leads to loss of the function. In the social amoeba *Dictyostelium discoideum*, obligate social cheaters cannot become dead stalk cells that lift spores up for dispersal, but instead depend on forming chimeras with fully functional altruistic individuals for forming a stalk. Obligate cheaters in *D. discoideum* are known to pay the cost of being unable to form fruiting bodies on their own. In this study we discovered that there are two additional costs that can apply to obligate cheaters. Even when there are wild-type cells to parasitize, the chimeric fruiting bodies that result have shorter stalks that are disadvantaged in spore dispersal. Furthermore, we found that obligate cheaters were overrepresented among spore cells in chimeras only when they were at low frequencies. Failure to develop into viable fruiting bodies on their own, negative frequency-dependent cheating, and shorter fruiting bodies represent three limits on obligate social cheating so it is not surprising that obligate cheaters have not been found in nature.

## Introduction

Cooperative behavior is common in nature, but cooperators are vulnerable to cheaters who can gain the benefits of cooperation without paying the costs [1]. In order for cooperation to persist, strong conflict between cooperators and cheaters must be mitigated [2–4]. Cheating can be reduced or eliminated by natural selection if the benefits of cooperation preferentially go to relatives because relatives are likely to share the gene or genes underlying the cooperative behavior. This is called kin selection, and is based on inclusive fitness theory, because individuals maximize their inclusive fitness, which includes their personal fitness as well as their effects on the fitness of their relatives, modified by how closely related they are [5–8].

The social amoeba *Dictyostelium discoideum* can form a multicellular fruiting body but requires altruistic action by a subset of cells in order to do so. In the wild, individual amoebas live in soil and leaf litter where they prey upon bacteria. When starved, they aggregate, develop into a multicellular slug, and migrate to a new location where they form a fruiting body. In this multicellular structure a minority of cells in the slug altruistically sacrifice their lives to form a dead stalk which lifts the other cells a few millimeters above the soil as viable, hardy spores in a structure called the sorus [9]. Altruism like this can evolve by kin selection if the spores are genetic relatives of stalk cells, provided the benefits of making a stalk are high enough.

Given that it is costly, why become stalk at all? The benefit of making a fruiting body seems to be dispersal, as those who have their stalks experimentally destroyed are dispersed less by a model insect vector *Drosophila melanogaster* than those with intact stalks [10].

Aggregations can form between unrelated genotypes [11] or even different species [12], setting the stage for social conflict between who becomes spore and who becomes stalk. In heterogeneous aggregations, called chimeras, natural selection should favor genotypes that preferentially become spores and place the burden of stalk-building on the other genotype. Conflict can be controlled in this system by high relatedness [13–15], as well as by other mechanisms such as pleiotropy [16] and a lottery-like role assignment system based on the cell cycle and nutrition [17]. In *D. discoideum*, the high relatedness necessary for preserving cooperation can be generated by active processes like kin discrimination [18–21], or by passive processes like spatial population growth and fine-scale population structure [22,23].

There are multiple ways for an amoeba to cheat [17,24–27]. Facultative cheaters will overrepresent themselves in the spores when in chimera but can still make fruiting bodies on their own. In *D. discoideum* the mutants *chtB*^*-*^ and *chtC*^*-*^ are examples of this strategy [28,29]. Fixed cheaters always allocate the same amount to spores and can be overrepresented in the spores if their fixed strategy happens to be to give more to spores than their social partner does. Allocation to spore vs stalk varies in nature so this may be common [30]. Social parasites, or obligate social cheaters, cannot make fruiting bodies on their own and tend to become spore in chimera such as the mutant *chtA*^*-*^, better known as *fbxA*^*-*^ [31]. Unlike the other categories of cheating, the spread of obligate social cheaters can threaten cooperation itself.

Relatedness in *D. discoideum* fruiting bodies is high in nature which keeps obligate social cheaters such as *fbxA*^*-*^ from spreading because they lack other genotypes to exploit [15]. Consistent with this idea, no obligate social cheaters have been isolated from nature despite extensive sampling [15,30]. In contrast, when relatedness is experimentally lowered in the lab, obligate social cheaters evolve readily and repeatedly [14,31,32]. However, resistors to cheating also evolve in these populations [33].

Aside from low fitness when alone and resistance evolution, there may be other potential limits to the spread of obligate social cheaters. One of these is negative frequency-dependent cheating. This occurs when cheaters are overrepresented in the spores only when they are at low frequencies. When cheating is negatively frequency-dependent, obligate social cheaters do not threaten cooperation itself because they are self-limiting. There is evidence for negative frequency-dependent cheating in *D. discoideum* for some facultative cheaters [26,34]. We know little about frequency-dependent cheating in obligate social cheaters. The only well-studied obligate cheater, *fbxA-*, overrepresents itself in the spores at all frequencies [15].

Another factor that could limit the spread of obligate social cheaters is if a cheater places the entire burden of stalk production on its social partner. If the partner does not compensate by allocating more to stalk than it would in a clonal fruiting body, then the fruiting body would be shorter. This may in turn reduce the likelihood that individuals in that sorus are dispersed as often or as far. The impact of social behavior on dispersal potential via the height of fruiting bodies, rather than their presence or absence, has not yet been experimentally tested.

Here we examine the effect of social conflict between a wild clone of *D. discoideum*, NC28.1, and an obligate, non-fruiting social cheater previously evolved in the laboratory from the same clone under conditions of low relatedness, called EC2 [14]. This obligate social cheater cannot develop properly on its own, similarly to *fbxA-*, and overrepresents itself in the spores when at a 10% initial frequency relative to NC28.1. We mixed these two clones at different frequencies and measured the heights of fruiting bodies they produced and the frequency of the obligate cheater in those fruiting bodies. We expected that fruiting bodies containing more obligate social cheaters would be shorter because the cheaters do not contribute to the stalk. Alternatively, the initial frequency of cheaters could have no effect on stalk height if the wild-type facultatively increases its allocation to stalk in order to increase the likelihood that it is dispersed, or the cheater coerces the wild-type to allocate to stalk, as has been found in some cheater mutants [26,35]. As far as frequency dependence goes, we predicted that another limitation to the success of obligate cheaters would be their decline in cheating as their proportion increases.

## Materials and Methods

### Strains and Culture conditions

To prepare food bacteria for *D. discoideum* clones to prey upon, we first spread non-pathogenic *K. pneumoniae* KpGe (Dicty Stock Center, dictybase.org) frozen in 80% KK2 [2.25 g KH_2_PO_4_ (Sigma-Aldrich) and 0.67 g K_2_HPO_4_ (Fisher Scientific) per liter] and 20% glycerol on an SM/5 agar media [2 g glucose (Fisher Scientific), 2 g yeast extract (Oxoid), 0.2 g MgCl_2_ (Fisher Scientific), 1.9 g KHPO_4_ (Sigma-Aldrich), 1 g K_2_HPO_5_ (Fisher Scientific), and 15 g agar (Fisher Scientific) per liter] and allowed the bacteria to grow at room temperature until single colonies appeared, which happened in about two days. We picked a single colony from this plate with a sterile loop, spread it on a new SM/5 plate, and allowed the bacteria to grow for two days in order to reach high abundance. We collected these bacteria into KK2 with a sterile loop and diluted them to 1.5 OD_600_ in KK2 (∼5 × 10^8^ cells, measured with an Eppendorf BioPhotometer). We used these bacteria as food for amoebas in our experiment and repeated this process anew for each of the three replicate experiments.

To grow NC28.1, the wild-type ancestor clone, from freezer stocks for use in our experiments we added spores frozen in 80% KK2 and 20% glycerol to 200*μ*l of 1.5 OD_600_ *K. pneumoniae* suspension. We spread the mix of spores and bacteria on SM/5 plates with a sterile glass spreader, then incubated the plates at room temperature for 7 days under constant overhead light until the social cycle was complete and fruiting bodies had formed. We repeated this process for each of the three replicate experiments.

To grow EC2, the RFP-labelled obligate social cheater [14], from freezer stocks for use in our experiments, we added amoebas frozen in HL5 (5 g proteose peptone, 5 g thiotone E peptone, 10 g glucose, 5 g yeast extract, 0.35 g Na2HPO4 * 7H2O, 0.35 g KH2PO4 per liter) with 10% DMSO to 200*μ*l of 1.5 OD_600_ *K. pneumoniae* suspension. We used amoebas rather than spores because EC2 produces few or no viable spores on its own. We spread the mix of amoebas and bacteria on an SM/5 plate with a sterile glass spreader, then incubated the plate at room temperature for 24-48 hours until starving EC2 amoebas began aggregating. We then used a sterile loop to transfer a sample to a new plate containing fresh *K. pneumoniae* for them to prey upon. These were allowed to grow for 24-48 hours until a vegetative front of amoebas had formed. We collected these amoebas with a sterile loop into ice-cold KK2 (see “Experimental procedures”). and ensured that the amoebas we used were clonal by plating 10 SM/5 plates with about 10 amoebas each, then picking a single clonal plaque originating from a single amoeba. We repeated this process for each of the three replicate experiments.

### Experimental procedures

In order to obtain cells of both *D. discoideum* clones for experimental mixing, we plated amoebas (EC2) or spores (NC28.1) previously grown from freezer stocks as described above on separate SM/5 agar plates with 200*μ*l of 1.5 OD_600_ *K. pneumoniae* suspension. We allowed the *D. discoideum* spores to proliferate until they reached high density, but before they aggregated and attempted to fruit, 24-48 hours.

We collected amoebas to make the mixtures by pouring ice-cold KK2 onto the plates, rubbing them into suspension with a gloved fingertip, then collecting and centrifuging the mixture at 10*°*C for 3 minutes at 1300 rpm in order to pellet the amoebas and leave *K. pneumoniae* in solution. We decanted the pellets, resuspended them in KK2, and measured their density with a hemacytometer before making the mixtures. For each treatment, we mixed 200*μ*l of fresh *K. pneumoniae* suspension with a total of 2×10^5^ amoebas then spread the solution evenly with an ethanol-sterilized glass spreader on an SM/5 agar plate. We made mixtures of EC2 and NC28.1 with various initial frequencies of EC2 (0.0, 0.1, 0.3, 0.5, 0.7, 0.9, and 1.0) in order to generate variation in their final frequencies in fruiting bodies. We repeated this experiment three times, each on a separate day.

We collected fruiting bodies after one week at room temperature under constant overhead light to allow fruiting bodies to fully develop. On each plate, we selected three fruiting bodies at random. To do this, we placed a plate of fruiting bodies over a grid of 1cm by 1cm squares with some of the squares colored in at random. We selected three of the colored-in squares at random using a random number generator and marked each plate at the centers of each of the squares. We then individually collected the closest fruiting body to each mark with fine tweezers. For each, we pressed the sorus, which contains the spores, against the side of a 100*μ*l tube containing 100*μ*l of KK2, to dislodge the spores then laid the stalk on a glass microscope slide. The contents of the tube were vortexed and immediately run through an Accuri C6 flow cytometer before they could stick to the sides of plastic tubes. We used 50*μ*l of suspended spores and recorded the numbers of fluorescent (EC2) and non-fluorescent (NC28.1) spores. After three fruiting bodies were collected from a single plate, the stalks were covered with a cover slip and sealed with nail polish for later imaging. Stalk length was individually recorded by imaging picked stalks under a Leica S8AP0 dissecting microscope with Leica application suite software v4.1 using the “draw line” tool.

### Analysis

We excluded several data points from the analysis for which we could not accurately measure stalk height due to damage during collection. We also excluded a data point for which very few spores were counted by the flow cytometer (<300) because the cheater proportion calculated for this sorus is probably inaccurate. We treated failed aggregates which did not fully differentiate into fruiting bodies as having a stalk height of zero (but we reanalyzed these data excluding failed aggregates and found similar results).

We had a treatment consisting entirely of cheaters (EC2). Though they generally do not produce fruiting bodies, they do succeed in making a very few. We did not include those data because our main question is whether mixtures result in shorter stalks. However, we did collect spores from these fruiting bodies in order to measure the proportion of EC2 cells which retained the RFP label through the process of fruiting. We use this to correct our proportion of fluorescent cells data from the other treatments to reflect the true proportion of EC2.

In order to test our whether increasing cheater frequency yields shorter fruiting bodies, we used a linear mixed-effects model with the function lme in the nlme package in R (version 4.2.1, The R Foundation for Statistical Computing) with stalk height as the response variable, the initial cheater frequency as a fixed effect, the total number of spores per sorus as a fixed effect, and the day of the experiment as a random effect (stalk height ∼ initial cheater frequency + total spores + 1|day). We included the total number of spores as a fixed effect in our initial model because fruiting bodies may be tall simply because they develop from larger populations.

We then excluded the random effect of day by using the lm function in base R and compared the two models with the anova function in base R (stalk height ∼ initial cheater frequency + total spores). We found that the two models were not significantly different (p = 0.27, with day: AIC = 105.67, without day: AIC = 104.89), so we proceeded with the simpler model without the effect of day. We then further simplified the model by removing the effect of total number of spores because it did not significantly affect stalk height (p = 0.20) (stalk height ∼ initial cheater frequency).

We also tested if the obligate social cheater clone, EC2, overrepresented itself in the spores of fruiting bodies relative to its initial frequency with a linear mixed-effects model using the function lme in the nlme package in R with final cheater proportion as the response variable, the initial cheater frequency as a fixed effect, and the day of the experiment as a random effect (final cheater proportion ∼ initial cheater proportion + 1|day). We excluded the random effect of day by using the lm function in base R and compared the two models with the anova function in base R (final cheater proportion ∼ initial cheater proportion). We found that the two models were not significantly different (p = 1, with day: AIC = −47.65, without day: AIC = −49.65, so we proceeded with the simpler model without the effect of day.

We further tested if the obligate social cheater clone, EC2, overrepresented itself in the spores of fruiting bodies at each initial frequency. We used one sample t-tests to test if the mean of the proportion of cheaters in fruiting bodies was significantly different from their initial frequency using the t.test() function in base R, setting mu to the initial cheater frequency.

Data and code for analysis are available at the Washington University Digital Research Materials Repository. DOI: https://doi.org/10.7936/exnd-2145

## Results

We mixed two clones of *D. discoideum* which were a wild-type ancestor and an obligate social cheater previously experimentally evolved from that ancestor, at various frequencies relative to one another and allowed them to form fruiting bodies together. We had two goals, to see the impact of increasing proportions of the social cheater on (1) stalk height and (2) on frequency of that cheater among the spores. We found that initial cheater proportion significantly predicted stalk height (linear model, DF = 48, t = −6.17, p = 1.37e-07). The slope of this relationship was negative (−1.34, SE = 0.22), indicating that increasing the initial cheater proportion decreased stalk height (Figure 1). We then reanalyzed the data excluding failed aggregates instead of treating them as having a stalk height of zero. We found that initial cheater proportion still significantly predicted stalk height (linear model, DF = 43, t = −4.83, p = 1.75e-05). The slope of this relationship remained negative (−1.22, SE = 0.25).

**Figure 1:**
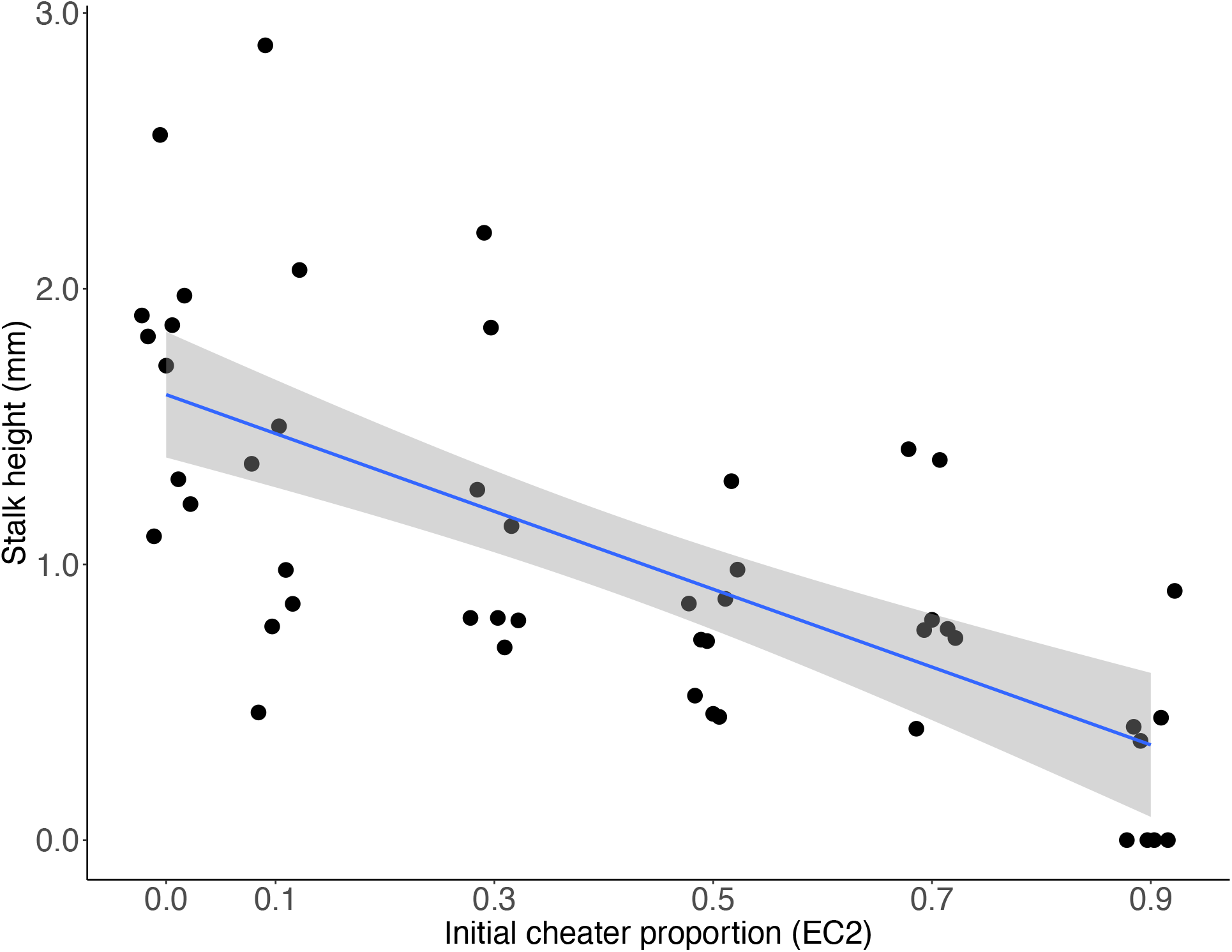
Initial cheater proportion predicts stalk height (linear model, DF = 48, t = −6.17, p = 1.37e-07). Regression line is y = −1.34x + 1.6. R-squared = 0.47. Shaded area is 95% CI of the slope of the line.

For the second goal on obligate social cheater representation in the spores relative to initial frequency, we found that initial cheater proportion significantly predicted final cheater proportion (linear model, DF = 38, t = 13.80, p = 2.23e-16). The slope of this relationship was positive and, most importantly for our question, significantly lower than 1 (0.85, SE = 0.06, p = 0.02), indicating the clone that did not make fruiting bodies on its own cheated its wildtype partner less as the frequency of the cheater increased (Figure 2).

**Figure 2:**
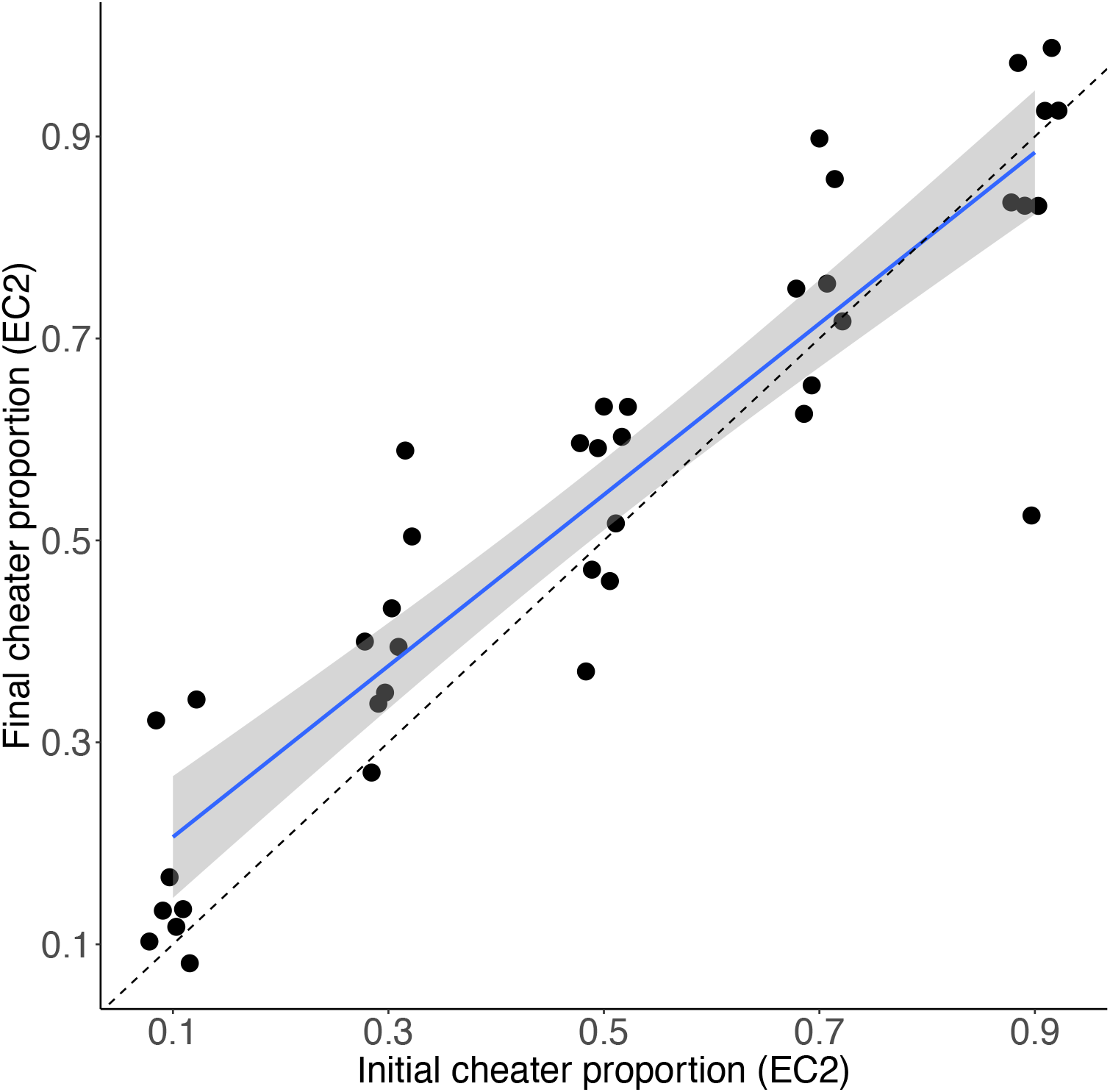
Initial cheater proportion predicts final cheater proportion (linear model, DF = 38, t = 13.80, p = 2.23e-16). Regression line is y = 0.85x + 0.12. R-squared = 0.83. Shaded area is 95% CI of the slope of the line. The dashed line shows the null hypothesis of a 1:1 relationship between initial cheater proportion and final cheater proportion.

When we tested each initial cheater frequency separately we found that cheating was evident only at lower frequencies of the cheating clone. At 0.1 initial frequency the final frequency was marginally greater than initial frequency (t = 2.1167, df = 7, p-value = 0.07207). At 0.3 initial frequency the final frequency was significantly greater than the initial frequency (t = 3.1008, df = 7, p-value = 0.0173). At 0.5, 0.7 and 0.9 initial frequencies the final frequency was not significantly different from the initial frequency (t = 1.3555, df = 8, p-value = 0.2123; t = 1.3493, df = 6, p-value = 0.2259; t = −1.3388, df = 8, p-value = 0.2174 respectively).

## Discussion

Social behavior like cooperation is common in microbes but populations of cooperators can be invaded by cheaters that benefit from cooperation without paying the cost [36,37]. Obligate social cheaters in particular can pose a threat to cooperation itself. Previous studies in *D. discoideum* found that an obligate social cheater mutant called *fbxA*^*-*^ is limited by poor spore production when it is without other clones to exploit [15,31]. We investigated a different obligate social cheater called EC2 for additional costs of this kind of social cheating.

First we found that chimeric fruiting bodies have shorter stalks. Shorter stalks could prevent cheaters from reaching high frequencies in a population because fruiting bodies with more obligate cheaters in them may be less likely to be picked up by an insect vector, which is the function of having a stalk (smith et al. 2014). Forming a fruiting body with a stalk appears to be important for *D. discoideum*’s fitness. Producing the fruiting body with its stalk is a complex process and is unlikely to be a side effect of some other selected trait. The importance of having a stalk is indicated by its presence across the entire ancient Dictyostelid family (though one small genus produces an acellular stalk) [38]. Finally, sacrificing some potential spores to produce a stalk can only be explained if there is some benefit. While we have not documented the fitness cost of shorter stalks, we know that a stalk height of zero can reduce dispersal [10] and it seems improbable that this disadvantage would not also apply also to at least very short stalks.

Cheaters can be either self-promoting, changing their own allocation to stalk, or coercive, forcing their partner to change its allocation. If the cheater is fully coercive, we would expect to see full-length stalks up to about 80% cheaters because the 20% wildtype has enough cells to make a normal stalk. Stalks would get smaller only after that point. Instead we see that increasing the frequency of obligate social cheater cells yields shorter fruiting bodies across the entire range. This is more consistent with the cheater being self-promoting, allocating more of its own cells to spores than to stalk.

In addition, we found that EC2, unlike *fbxA*^*-*^, overrepresented themselves in the spores only when they were mixed at low frequencies. This negative frequency dependence should help prevent EC2 from becoming common and causing the complete breakdown of cooperation. There is some evidence for frequency-dependent cheating in facultative cheaters in *D. discoideum* such that cheating is weaker at higher frequencies [26,34] but not for the obligate cheater *fbxA*^*-*^*[15]*. Frequency-dependence may be a common limitation on cheating in *D. discoideum*, but the reasons why cheaters do worse at higher frequencies are unclear. It could be that the benefit of cheating decreases as cheaters increase in frequency because there are fewer cooperators to exploit. Frequency-dependent cheating is one way that cooperation can be maintained despite the presence of obligate cheaters and has been found in other microbes such as *P. aeruginosa* [39], *M. xanthus* [40], and *Bacillus subtilis* [41].

Cheating can occur whenever the benefits of cooperation can be exploited by those that do not pay the cost of cooperating. However, it is important to understand natural population structure when evaluating the likelihood that a cheating mutation will spread in nature. In this example, obligate social cheaters like these would be unlikely to spread to fixation since relatedness in nature is typically high [15] due to both limited dispersal [22,23]and active kin discrimination [18]. This means that obligate cheaters should they arise in nature will aggregate and develop with clonemates and therefore be unable to fully differentiate into fruiting bodies. In addition, no nonfruiting mutants of *D. discoideum* have ever been found in nature, even though they can be easily evolved in the lab under conditions of low relatedness [14,15,30,32]. The disadvantages when clonal of not forming spores at all and when not clonal of forming short fruiting bodies are simply too great.

High relatedness is also expected to impact the two new limits on cheating we have observed. At low relatedness, groups will be genetically well mixed and so stalk heights will be uniform, leading to little selection against short-stalked cheaters. Under high relatedness, cheaters and cooperators will be mostly separated, leading to tall cooperator fruiting bodies and short cheater fruiting bodies. The frequency-dependent disadvantage is a little more complicated. At low relatedness, each fruiting body will match the population frequency of cheaters, and cheaters will spread up to the equilibrium frequency where they have no more advantage. At high relatedness, there will always be variation, with cheaters getting an advantage in fruiting bodies with few cheater cells, and no advantage in fruiting bodies with many cheaters. High relatedness may actually therefore weaken frequency-dependent control of this cheater because, at or above the low-relatedness equilibrium frequency, there will still be some groups that can cheat. However this might change for a cheater that was actually disadvantageous rather than neutral at high frequency.

Similar issues of cheating and relatedness occur in other systems. For example, different strains of the bacterium *Myxoccocus xanthus* can exhibit strong antagonism against one another [42]. However, natural populations of *M. xanthus* are highly structured so mixing between genotypes is unlikely [43]. Furthermore, kin discrimination that segregates genotypes evolves rapidly in this species under laboratory conditions [44]. The costs of cheating to groups that they are a part of is called cheating load, which can be substantial [24]. Similarly to obligate social cheaters in *D. discoideum*, obligate social cheaters in *M. xanthus* can lead to population collapse when at high frequency [45].

Our results provide evidence for two mechanisms other than failed spore production that could prevent the spread of obligate social cheaters in *D. discoideum*. The first is reduced potential for dispersal and the second is when cheating is frequency-dependent. This is interesting because EC2 has evolved from a wild ancestor while *fbxA*-is an artificial mutant created in a lab-adapted background strain. For this reason cheating in EC2 may reflect cheating in nature more closely than cheating in *fbxA*-. These additional limits on obligate social cheating in *D. discoideum* paint a more complete picture of why obligate cheaters do not spread in nature and cause the collapse of cooperation.

## Acknowledgements

Funding provided by National Science Foundation IOS 16-56756 and DEB 17-53743.

